# Ketamine-Induced Unresponsiveness Shows a Harmonic Shift from Global to Localised Functional Organisation

**DOI:** 10.1101/2024.06.20.599885

**Authors:** M. Van Maldegem, J. Vohryzek, S. Atasoy, N. Alnagger, P. Cardone, V. Bonhomme, A. Vanhaudenhuyse, A. Demertzi, O. Jaquet, M. A. Bahri, M. L. Kringelbach, E. A. Stamatakis, A. I. Luppi

## Abstract

Ketamine is classified as a dissociative anaesthetic that, in sub-anaesthetic doses, can produce an altered state of consciousness characterised by dissociative symptoms, visual and auditory hallucinations, and perceptual distortions. Given the anaesthetic-like and psychedelic-like nature of this compound, it is expected to have different effects on brain dynamics in anaesthetic doses than in low, sub-anaesthetic doses. We investigated this question using connectome harmonic decomposition (CHD), a recently developed method to decompose brain activity in terms of the network organisation of the underlying human structural connectome. Previous research using this method has revealed connectome harmonic signatures of consciousness and responsiveness, with increased influence of global network structure in disorders of consciousness and propofol-induced sedation, and increased influence of localised patterns under the influence of classic psychedelics and sub-anaesthetic doses of ketamine, as compared to normal wakefulness. When we applied the CHD analytical framework to resting-state fMRI data of volunteers during ketamine-induced unresponsiveness, we found increased prevalence of localised harmonics, reminiscent of altered states of consciousness. This is different from traditional GABAergic sedation, where instead the prevalence of global rather than localised harmonics seems to increase with higher doses. In addition, we found that ketamine’s harmonic signature shows higher alignment with those seen in LSD- or psilocybin-induced psychedelic states than those seen in unconscious individuals, whether due to propofol sedation or brain injury. Together, the results indicate that ketamine-induced unresponsiveness, which does not necessarily suppress conscious experience, seems to influence the prevalence of connectome harmonics in the opposite way compared to GABAergic hypnotics. We conclude that the CHD framework offers the possibility to track alterations in conscious awareness (e.g., dreams, sensations) rather than behavioural responsiveness – a discovery made possible by ketamine’s unique property of decoupling these two facets.

## Introduction

In 1966, Corssen and Domino published the first clinical study investigating ketamine as a human anaesthetic. They discovered that ketamine could swiftly induce strong pain relief alongside a unique state of altered consciousness, and that the limited duration of effect could be safely extended with repeated administration (Corssen & Domino, 1966). Later studies found that increasing concentrations, ranging from 0.3 to 1 mg/kg/hour, initially result in antidepressant action, followed by pain relief and altered perception, such as sensations of detachment, visual distortions, and confused thinking (Berman et al., 2000; Krystal et al., 1994; Lavender, Hirasawa-Fujita & Domino, 2020; Olofsen et al., 2022; Peltoniemi et al., 2016). Loss of responsiveness is usually only observed with the administration of high doses, ranging from 1 to 6 mg/kg/hour (Barrett et al., 2020; Lavender, Hirasawa-Fujita & Domino, 2020; Sleigh et al., 2014). Due to these widespread pharmaceutical effects, ketamine stands out as an incredibly versatile medication utilised for anaesthesia and various other medical purposes, such as treatment-resistant depression and pain relief (Abdallah et al., 2022; Johnston et al., 2024; Kurdi, Theerth & Deva, 2014; Nowacka & Borczyk, 2019).

So far, a substantial body of research has explored the neurophysiological effects of ketamine-induced unresponsiveness (Akeju et al., 2016; Blain-Moraes et al., 2014; Bonhomme et al., 2016; Colombo et al., 2019; Lee et al., 2013; Li, Vlisides & Mashour, 2022; Mashour, 2016; McMillan & Muthukumaraswamy, 2020; Varley et al., 2020b, 2021; Vlisides et al., 2017). At the molecular level, ketamine acts primarily as a non-competitive antagonist of the *N*-methyl-*D*-aspartate (NMDA) receptor by blocking its channels and decreasing their opening frequency (Anis et al., 1983; Bergman, 1999; Domino, Chodoff & Corssen, 1965; Hirota & Lambert, 2022; Orser, Pennefather & MacDonald, 1997; Sattar et al., 2018; Sleigh et al., 2014; Thomson, West & Lodge, 1985). This results primarily in suppression of GABAergic inhibitory cortical interneurons, generating an overall increase in excitability throughout the brain (Homayoun & Moghaddam, 2007; Seamans, 2008). For a thorough discussion of the functioning of NMDA receptors (Dupuis, Nicole & Groc, 2023; Ogden & Traynelis, 2011; Paoletti, Bellone & Zhou, 2013; Vance et al., 2011) and ketamine’s interaction with other molecular systems (Li & Vlisides, 2016), we refer the reader to the relevant literature.

At the systems neuroscience level, the mechanisms of ketamine sedation show major differences from those of more traditional anaesthetics, such as propofol and sevoflurane. For example, ketamine seems to suppress sleep-promoting regions of the hypothalamus, activate arousal-promoting regions of the brainstem and diencephalon, preserves cross-modal interactions between sensory regions, and increase EEG and MEG gamma activity in the cortex (Akeju et al., 2016; de la Salle et al., 2016; Ferrer-Allado et al., 1973; Lü et al., 2008; Mashour, 2014, 2016; Mashour & Hudetz, 2017; Pal et al., 2015; Shaw et al., 2015), while GABAergic anaesthetics induce opposite changes (Murphy et al., 2011; Yue et al., 2021). In addition, ketamine is thought to interrupt information transfer between cortical regions, while still maintaining the function of sensory networks and subcortical areas (Blain-Morraes et al. 2014; Joules et al., 2015; Lee et al., 2013; Schroeder et al., 2016). Despite some similarities with more traditional anaesthetics (Lee et al., 2013; Wang et al., 2017), ketamine thus seems to induce a specific large-scale organisation of functional connectivity throughout the brain, which might be associated with its distinct molecular targets (Adam et al., 2024).

Another unique characteristic of ketamine-induced unresponsiveness is its ability to preserve some conscious experiences, most often seen in the form of vivid dreams and sensations (Collier, 1972; Bonhomme et al., 2016; Garfield et al., 1972; Hejja & Galloon, 1975; Krestow, 1974; Maya et al., 2018; Noreika et al., 2011; Sarasso et al., 2015). This is a useful property of the compound, as it allows researchers to disentangle the concepts of behavioural responsiveness and conscious awareness (Boly et al., 2013). Until recently, the ability to provide coherent responses to external stimuli served as the primary criterion for assessing consciousness of non-communicating patients and non-human animals (Chernik et al., 1990; Giacino, Kalmar & Whyte, 2004; Gao & Calderon, 2020). Yet, this method operates under the assumption that unresponsive states uniformly signify unconsciousness, despite evidence indicating that vivid experiences can manifest within such states. For example, dreams have been observed during various stages of sleep (Nielsen, 2000; Solms, 2000; Siclari et al., 2017, 2018) and even under general anaesthesia (Noreika et al., 2011). Moreover, after severe brain injury, some patients remain unresponsive at the bedside yet show some level of sensorily disconnected consciousness through neuroimaging (Thibaut et al., 2021; Vanhaudenhuyse et al., 2018). Conversely, some patients have episodes of connected consciousness during surgical procedures under general anaesthesia where they may experience pain and awareness despite appearing unresponsive (Ghoneim et al., 2009; Sanders et al., 2017). It seems thus crucial to differentiate between the brain dynamics associated with consciousness and those related to behavioural responsiveness (Arena et al., 2022).

Multiple lines of evidence have shown that human consciousness relies on a dynamic repertoire of brain activity (Barttfeld et al., 2015; Campbell et al., 2020; Demertzi et al., 2019; Golkowski et al., 2019; Gutierrez-Barragan et al., 2021; Huang et al., 2016, 2020; Lord et al., 2023; Luppi et al., 2019, 2021b; Noirhomme et al., 2010; Tanabe et al., 2020), which prompts the inquiry into how a static network of anatomical connections, the human connectome, can give rise to these complex dynamics (Abdelnour, Voss & Raj, 2014; Atasoy et al., 2018a; Deco, Jirsa & McIntosh, 2011; Cabral, Kringelbach & Deco, 2017; Fukushima et al., 2018; Xie et al., 2021). Early exploration down these lines of enquiry has developed our understanding of the loss of consciousness and its signatures: as consciousness fades, the patterns of correlation among regional BOLD timeseries (functional connectivity) tend to mirror more closely the pattern of anatomical connections between these regions (Barttfeld et al., 2015; Demertzi et al., 2019; Gutierrez-Barragan et al., 2021; Lee et al., 2019; Ma, Hamilton & Zhang, 2017; Tagliazucchi et al., 2016a, Ulrig et al., 2018). In contrast, brain activity patterns tend to be more localised in altered states of consciousness, such as psychedelic and meditative states (Atasoy et al., 2017, 2018b, 2023; Luppi et al., 2023b; Tagliazucchi et al., 2014; Varley et al., 2020a; Vohryzek et al., 2024).

One way to investigate the functional organisation of brain activity is through the mathematical framework of connectome harmonic decomposition (CHD; Atasoy et al., 2016). In CHD, the well-known Fourier transform, which decomposes a signal in the time domain (complex wavefunction) into a set of basic functions (basic sinusoidal waves) known as temporal harmonics (Figure 1A), is extended to the network architecture of the human brain (Figure 1B). Functional brain signals undergo a transformation from the spatial domain to the domain of connectome harmonics, where they manifest as distributed activity patterns, each linked to a distinct spatial frequency ranging from broad to intricate (Figure 1B; Atasoy, Donnelly & Pearson, 2016; Atasoy et al., 2017, 2018a; Brahim & Farrugia, 2020). Functional MRI signals can then be decomposed into these basic components (connectome harmonics; Figure 1C-G), each with their own contribution to the original signal. An increased contribution of low-frequency (coarse-grained) harmonics indicates more global brain activity patterns, while an increased contribution of high-frequency (fine-grained) harmonics corresponds to more localised patterns. A detailed account of the procedure can be found in the ‘Methods’ section.

**Figure 1.**
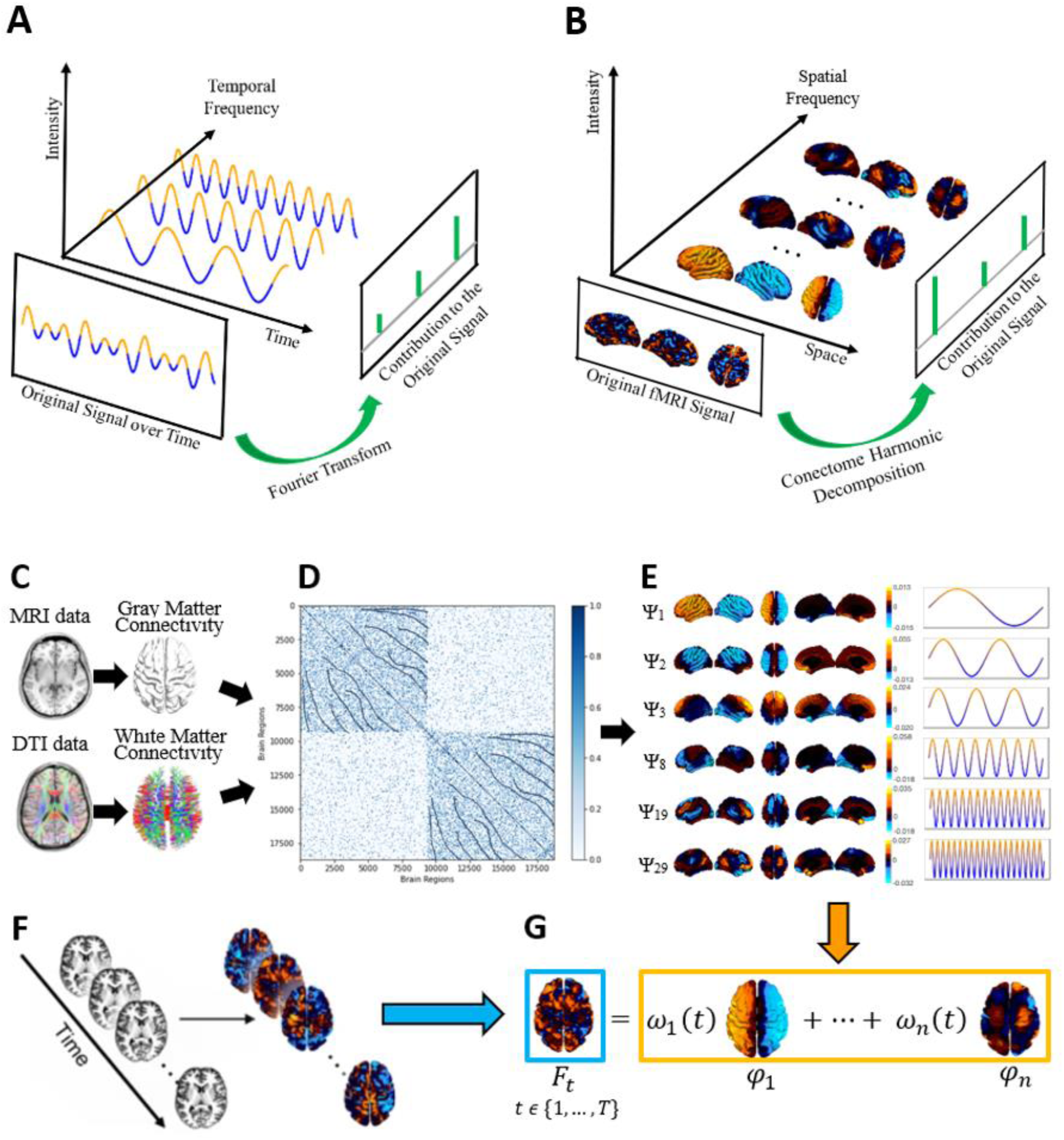
Connectome Harmonic Decomposition of Resting-State fMRI Data. **A** Traditional Fourier analysis decomposes a time-series signal (e.g. complex wave function) into temporal harmonics of different frequency (basis wave functions). Temporal harmonics with high frequencies represent rapidly changing signals, where data points close in time can have different values. On the other hand, low-frequency temporal harmonics correspond to signals that change slowly over time, resulting in temporally adjacent data points having similar values, indicating a stronger time-dependent nature of the signal. **B** Connectome harmonic decomposition decomposes a signal originating in the spatial domain (expressed as BOLD activation across distinct spatial points throughout the cortex) into harmonic modes of the human structural connectome. This process yields a new set of basic functions characterised by widespread patterns of activity propagation across the brain, operating at various scales. **C** T1 weighted magnetic resonance imaging (MRI) data is employed to reconstruct the cortical surface between grey and white matter. Diffusion tensor imaging (DTI) data is utilised to extract thalamo-cortical fibers through deterministic fiber tractography. **D** Local and long-distance connections are integrated to construct the connectivity matrix of the human connectome. **E** The graph Laplacian of this high-resolution connectome is subsequently broken down into its eigenvectors *φ*_1…*n*_ (harmonic modes) and their corresponding eigenvalues *λ*_1…*n*_ (spatial frequencies with ascending granularity) by solving the following equation: Δ_*G*_*φ*_*k*_(*v*_*i*_) = *λ*_*k*_*φ*_*k*_(*v*_*i*_). As the connectome harmonic number k (also referred to as wavenumber) increases, increasingly intricate and detailed spatial patterns are observed. **F** At each timepoint *t ∈* {*1, …, T*}, functional magnetic resonance imaging (fMRI) data is mapped from volumetric space onto the cortical surface. **G** Connectome harmonic decomposition (CHD) of the fMRI data assesses the contribution of *ω_k_*(*t*) of each harmonic mode *φ_k_* to the cortical activity at each timepoint *t ∈* {*1, …, T*}.

Luppi and colleagues (2023b) have previously suggested that the harmonic signature obtained by CHD reliably tracks alterations of consciousness, with an increased contribution of global harmonics in propofol sedation and disorders of consciousness (DOC) and an increased contribution of localised patterns in typical (LSD & psilocybin) and atypical (low doses of ketamine) psychedelic states. To make sure that they were not tracking mere behavioural responsiveness, the researchers used a dataset of patients (*N* = 22) that met the diagnostic criteria for disorders of consciousness, of which some (*N* = 8) demonstrated conscious awareness by successfully completing mental imagery tasks while in the MRI scanner (Owen et al., 2006). As expected, the results showed an increased contribution of global patterns in patients that did not show signs of consciousness compared to those that did (Luppi et al., 2023b). However, it is difficult to know with absolute certainty that the unresponsive patients had not preserved any form of consciousness, as some of them may have been conscious but unable to do the task for a variety of reasons (e.g., impairments in language processing, hearing difficulties).

Ketamine deep sedation has the unique property of temporarily suppressing behavioural responsiveness while preserving some form of conscious awareness (Bonhomme et al., 2016; Collier, 1972; Noreika et al., 2011; Sarasso et al., 2015). In contrast to behaviourally unresponsive DOC-patients, the pharmacological effects of ketamine manifest themselves only temporary, making it possible for participants to report experiences – if any – they had during the sedated period. The dataset used in this study provides such (anecdotal) reports – obtained through a phone call after completion of the study – showing that participants preserved some sort of consciousness even though they were behaviourally unresponsive (Bonhomme et al., 2016). By using the novel framework CHD on this special dataset, we aimed to build on the results of Luppi et al. (2023b) and further disentangle the concepts of behavioural responsiveness and conscious awareness in the neuroscientific study of consciousness.

Specifically, this study tries to clear out which of two opposite hypotheses is true. First, the connectome harmonic signature obtained by CHD could track behavioural responsiveness. In that case, brain activity during ketamine-induced unresponsiveness should show an increased contribution of global harmonics as compared to wakefulness. In contrast, the connectome harmonic signature could orchestrate consciousness, in which case we should see a similar or increased contribution of more localised patterns in ketamine-induced unresponsiveness as compared to wakefulness. To further statistically assess this, we will investigate the alignment between the harmonic signature of ketamine deep sedation and earlier identified signatures of other states of consciousness (propofol sedation, disorders of consciousness, LSD, ketamine sub-anaesthesia; Luppi et al., 2023b).

## Methods

### Subjects

The ketamine data used here were collected at the University Hospital of Liege (Liege, Belgium), registered at EudraCT 2010-023016-13, and have been published before (Bonhomme et al., 2016). For details, readers are encouraged to consult previous publications with these data (Bonhomme et al., 2016; Luppi et al., 2023a). In short, 14 right-handed volunteers (age 19 to 31, 5 women) were recruited through advertisement in an Internet forum and underwent medical interview and physical examination before their participation. Furthermore, every volunteer was provided with financial compensation to account for the inconvenience and time lost during the experiment (Bonhomme, 2016).

### Study Protocol

For a more detailed description of the study protocol, we direct the reader to the original publication with this data (Bonhomme et al., 2016). In short, the volunteers were instructed to refrain from consuming solids for a minimum of 6 hours and liquids for 2 hours prior to the experimental session. Upon entering the investigation unit, a thorough review of potential contraindications to participation was conducted, including criteria for both anaesthesia and MRI, using a comprehensive checklist. After structural MRI acquisition, subjects were removed from the scanner, and 64 electroencephalogram (EEG) scalp electrodes were placed to allow for simultaneous EEG recording during fMRI data acquisition (Brain Amp® magnetic resonance compatible acquisition setup; Brain Products GmbH, Germany). In this report, data analysis has been limited to fMRI data only.

Ketamine was delivered intravenously through a computer-controlled infusion device, which consisted of a distinct laptop computer connected to an infusion pump. The pharmacokinetic model employed to regulate the pump was the Domino model (Domino et al., 1982), which has been validated and shown to exhibit satisfactory predictive performance (Absalom et al., 2007). For each alteration in ketamine concentration, a 5-minute equilibration period was allowed upon reaching the target, to facilitate the equalisation of ketamine concentration across body compartments (Bonhomme et al., 2016).

The Ramsay Scale (RS; Ramsay et al., 1974) and the University of Michigan Sedation Scale (UMSS; Malviya et al., 2002) were used to assess the depth of sedation. Evaluation occurred directly before and after each fMRI data acquisition sequence. Volunteers were instructed to firmly squeeze the hand of the investigator, with the command reiterated twice. To ensure close monitoring of the volunteer, an investigator remained inside the MRI scan room throughout the process.

A first fMRI data acquisition was performed in the absence of any infusion of ketamine. Infusion was then started, and its target concentration was increased by steps of 0.5 μg ml^−1^ until a level of sedation corresponding to RS 3 to 4 or UMSS 1 to 2 was reached (light sedation, S1). After the 5-min equilibration period, a novel sequence of data acquisition occurred, consisting of the same sequence of events as during wakefulness. Ketamine target concentration was then further increased by steps of 0.5 μg/ml until RS 5 to 6 or UMSS 4 (deep sedation, S2), and the same sequence of data acquisition was again performed. Because ketamine has a long elimination half-life and to limit time spent in the fMRI scanner for the volunteer, the temporal order of those clinical states was not randomized. For the same reason, a recovery experimental condition could not be achieved. After those acquisitions, the infusion of ketamine was stopped, and the subject was removed from the fMRI scanner to allow for comfortable recovery. The presence of dreaming during ketamine infusion was checked through a phone call at distance from the experimental session (Bonhomme et al., 2016).

### MRI Data Acquisition

The acquisition procedures are described in detail in the original study (Bonhomme et al., 2016). Briefly, MRI data were acquired using a 3T Siemens Allegra scanner (Siemens AG, Germany; Echo Planar Imaging sequence using 32 slices; repetition time = 2460 ms; echo time = 40 ms; field of view = 220 mms; voxel size = 3.45 × 3.45 × 3 mm; matrix size = 64 × 64 × 32). For each volunteer in each condition, 300 functional volumes were acquired. A high-resolution structural T1 image was obtained in each participant at the beginning of the experiment for co-registration to the functional data. 6 participants had to be excluded from the study and further data analysis because of excessive agitation and movements (5 subjects) or voluntary withdrawal (1 subject), leaving 8 (Bonhomme et al., 2016).

### fMRI Preprocessing and Denoising

Preprocessing and denoising followed the same pipelines as in related previous publications (Luppi et al., 2019; Luppi et al., 2021b; Luppi et al., 2023b). For clarity and consistency of reporting, the same wording is used as in those publications.

“The functional imaging data were preprocessed using a standard pipeline implemented within the SPM12-based (http://www.fil.ion.ucl.ac.uk/spm) toolbox CONN (http://www.nitrc.org/projects/conn), version 17f (Whitfield-Gabrieli & Nieto-Castanon, 2012). The pipeline consisted of following steps: removal of the first five scans, to allow magnetisation to reach steady state; functional realignment and motion correction; slice-timing correction to account for differences in time of acquisition between slices; identification of outlier scans for subsequent regression by means of the quality assurance/artifact rejection software *art* (http://nitrc.org/projects/artifact_detect); structure-function co-registration using each participant’s high-resolution T1-weighted image; spatial normalisation to Montreal Neurological Institute (MNI-152) standard space with 2 mm isotropic resampling resolution, using the segmented grey matter image, together with an a priori grey matter template.

To reduce noise due to cardiac and motion artefacts, the anatomical CompCor method of denoising was used, also implemented within the CONN toolbox (Behzadi et al., 2007). Linear detrending was also applied, and the subject-specific denoised BOLD signal timeseries were band-pass filtered to eliminate both low-frequency drift effects and high-frequency noise, thus retaining temporal frequencies between 0.008 and 0.09 Hz. Importantly, this band-pass filtering pertains to temporal frequencies, which are distinct from the spatial frequencies obtained from connectome harmonic decomposition (as described below).” (Luppi et al., 2023b).

### Connectome Reconstruction

The following workflow is the same as described in previous work by Atasoy and colleagues (Atasoy et al., 2017). A high-resolution human structural connectome was obtained from diffusion tensor imaging (DTI) and structural MRI data from an independent sample of 10 human connectome project (HCP) subjects (6 female, age 23 to 35 years). For each subject, the cortical surfaces of each hemisphere at the interface of white and grey matter were reconstructed using Freesurfer (http://freesurfer.net), which resulted in a representation of 18,715 cortical surface vertices per participant. When each subject’s diffusion imaging and cortical surface data were co-registered, every vertex of the reconstructed cortical surface served as a centre to initialise eight seeds for deterministic tractography, performed using the MrDiffusion tool (http://white.stanford.edu/newlm/index.php/MrDiffusion). Tracking was terminated when fractional anisotropy (FA) was below a threshold of 0.3, with a minimum tract length of 20 mm, and a maximum angle of 30 degrees between consecutive tracking steps (Atasoy et al., 2017).

The structural connectivity of each participant was represented as a binary adjacency matrix, denoted as A, with each cortical surface vertex considered a node. For every pair of nodes *i* and *j* out of the *n* = 18,715 total nodes, *A_ij_* was set to 1 if a white matter tract connected them, as estimated from the deterministic tractography step described above, or if they were adjacent in the grey matter cortical surface representation. If neither long-range nor short-range connections between *i* and *j* existed, *A_ij_* was set to 0. The whole process resulted in a symmetric (undirected) binary matrix, as described earlier by Atasoy and colleagues (2017). Subsequently, the individual adjacency matrices of the 10 HCP participants were averaged to derive a group-average matrix *A̅*, which represents a typical structural connectome. Finally, the degree matrix *D* of the graph was defined, where each element *D*_*ij*_ represents the sum of connections for node *i* across all nodes 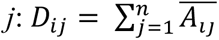 (Atasoy et al., 2017).

### Connectome Harmonics Extraction

Following the procedure used by Atasoy et al. (2017), the graph Laplacian Δ_*G*_ was computed on the group-average adjacency matrix *A̅*, described above, in order to estimate the discrete counterpart of the Laplace operator Δ (Chung, 1997).

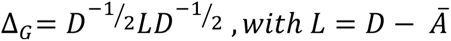

Subsequently, the connectome harmonics *φ*_*k*_, *k* ∈ {1, …, 18,715} were calculated by solving the following eigenvalue problem:

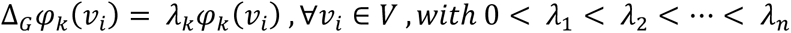

where *λ*_*k*_, k ∈ {1, …, *n*} is the corresponding eigenvalue of the eigenfunction *φ*_*k*_, *V* is the set of cortical surface vertices and *n* represents the number of vertices. The frequencies associated with each connectome harmonic are in the spatial rather than the temporal domain, and should not be confused with the temporal frequencies identified by Fourier transform in the temporal domain (e.g., timeseries denoising).

### Connectome-harmonic Decomposition of fMRI Data

At each timepoint *t* ∈ {1, …, *T*}, which corresponds to one TR, the pre-processed and denoised fMRI data were projected onto cortical surface coordinates by means of the HCP Workbench *-volume-to-surface-mapping* tool (Atasoy et al., 2017). Subsequently, the spatial pattern of cortical activity over vertices *v* at time *t*, denoted as *F_t_(v)*, was decomposed as a linear combination of the set of connectome harmonics 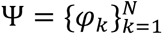:

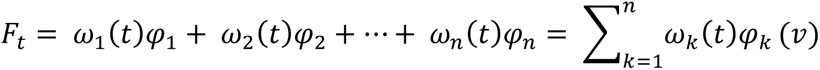

with the contribution *ω*_*k*_(*t*) of each connectome harmonic *φ*_*k*_ at time *t* being estimated as the projection (dot product) of the fMRI data *Ft(v)* unto *φ*_*k*_: *ω*_*k*_(*t*) = 〈*F*_*t*_, *φ*_*k*_〉 (Atasoy et al., 2017).

The transition from low-frequency (coarse-grained) to high-frequency (fine-grained) connectome harmonics indicates a move from global to more localised activity patterns (Atasoy et al., 2016, 2017). Other authors have interpreted the same shift as a departure of functional activity from the structural connectivity beneath it (Lioi et al., 2021; Medaglia et al., 2018; Preti & Van De Ville, 2019).

### Connectome Harmonic Power and Energy

Still following the procedure of Atasoy and colleagues (2017), the magnitude of contribution of each harmonic *φ*_*k*_, k ∈ {1, …, *n*} at any given timepoint *t*, called its “power”, is computed as the amplitude of its contribution: *P(φ*_*k*_, *t) = |ω*_*k*_(*t*)|.

In turn, the normalised frequency-specific contribution of each harmonic *φ*_*k*_, k ∈ {1, …, *n*} at timepoint *t*, called “energy”, is estimated by combining the strength of activation (|*ω*_*k*_(*t*)*|^2^*) of a particular connectome harmonic with its own intrinsic energy (*λ*_*k*_^2^).

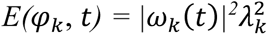

Consequently, total brain energy at time *t* is given by

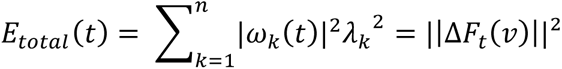

Finally, a binned energy spectrum across subjects and timepoints is construed by discretising the energy of connectome harmonics into 15 logarithmically-spaced frequency-specific bins, following previous work showing that this procedure can successfully highlight the connectome harmonic signatures of altered states of consciousness (Atasoy et al., 2017, 2018; Luppi et al., 2023b).

### Data-driven Extraction of Multivariate Connectome Harmonic Signatures

Partial Least Squares (PLS), also recognised as Projection on Latent Structures, is a multivariate statistical method used for modelling the relationship between one or multiple target variables (Y) and a set of predictor variables (X). The main goal is to find a set of latent variables, also known as principal components, that capture the maximal covariance with each other. In this study, X was the matrix of 15 binned energy values for each subject (averaged over timepoints), and Y was the vector of binary classification between the two states (awake vs. S1 and awake vs. S2). Due to the binary nature of Y, this analysis can be seen as an application of Partial Least Squares Discriminant Analysis (PLS-DA; Lee, Liong & Jemain, 2018). The first principal component represents the most discriminative pattern present in the data in terms of distinguishing between two states, which we term the state’s multivariate signature (MVS). This procedure has been introduced and described before (Luppi et al., 2023b).

### Diversity of Connectome Harmonic Repertoire

A diverse repertoire of connectome harmonics can be defined as a repertoire in which different harmonic modes contribute in different degrees to brain activity (Luppi et al., 2023b). This means that there is neither one dominating mode, corresponding to a periodic oscillation, nor equal contribution of every mode to the signal, corresponding to white noise. To quantify the connectome harmonic diversity, the entropy of the distribution of connectome harmonic power (absolute strength of contribution to the cortical activation pattern) across all 18,715 harmonics was calculated (for each timepoint of each subject). To deal with continuous data, the Kozachenko approximation was used, as implemented in the Java Information Dynamics Toolbox (JIDT; http://jlizier.github.io/jidt/) (Lizier et al., 2014). It is worth noting that when dealing with continuous variables, entropy can adopt negative values (Cover & Thomas, 2005). However, the interpretations remain the same: a more entropic distribution will correspond to a more diverse connectome harmonic repertoire.

### Statistics and Reproducibility

To be consistent with previous work in the field (Luppi et al., 2023b), statistical significance of the differences between conditions (wakefulness vs ketamine sedation) were assessed with Linear Mixed Effects models (MATLAB function: *fitlme*), considering condition as a fixed effect, and subjects as random effects. Timepoints were also included as random effects, nested within subjects, when a measure of interest was measured for each of those. Upper and lower bounds of the 95% confidence interval of the fixed effect, with associated *p*-value, are reported. We included demographic information (i.e., age and biological sex) as covariates of no interest. The False Discovery Rate for multiple comparisons across 15 frequency bins of harmonic energy was controlled by means of the Benjamini-Hochberg (Benjamini & Hochberg, 1995). Spearman’s non-parametric rank-based *ρ* was used for correlation. All analyses were two-sided, with an alpha threshold of 0.05.

## Results

In the current study, we utilise the CHD framework for the first time to extract the harmonic signatures underlying ketamine-induced unresponsiveness. In the subsequent section, we mainly report comparisons between the deepest stage of ketamine sedation (S2) and wakefulness (Awake), as these will be sufficient to test our hypotheses. However, comparisons between the lighter stage of ketamine sedation (S1) and wakefulness (Awake), and between the two different ketamine stages (S1 and S2) can be found in the Supplementary Materials and are briefly mentioned in the manuscript.

To substantiate our results, we applied the same analysis using an alternative reconstruction of the human connectome at higher spatial resolution. This reconstruction was obtained by aggregating data from 985 subjects from the HCP, arguably providing one of the most comprehensive representations of the human structural connectome currently available (Atasoy et al., 2017; Luppi et al., 2023b).

### Ketamine-Induced Unresponsiveness Increases Energy of Brain Activity

Initially, the activation power of each connectome harmonic was assessed by evaluating its contribution to the cortical activity pattern of the fMRI volume at each time point. Subsequently, by merging this activation power with the inherent, frequency-dependent energy of each connectome harmonic, its total energy was computed.

Utilising these fundamental metrics, obtained by connectome harmonic decomposition (CHD) of fMRI data, the overall impact of ketamine-induced unresponsiveness (deep sedation, S2) was investigated across the entire range of connectome harmonics. The analysis showed that the total energy of all these harmonics averaged across all time points significantly increases under ketamine deep sedation (S2) compared to normal wakefulness (*p* = 0.0257, repeated-measures t-test; Figure S1). Furthermore, the increase was also seen in lighter ketamine sedation (S1) as compared to wakefulness (*p* = 0.0146, repeated-measures t-test; Figure S1) and, although not significant, between S1 and S2 (*p* = 0.0788, repeated-measures t-test; Figure S1).

### Ketamine-induced Unresponsiveness is Characterised by an Increased Contribution of Localised Harmonics

Previous research using CHD has indicated that a sub-anaesthetic dose of ketamine, as compared to placebo, results in a decreased contribution of global (coarse-grained) connectome harmonics and an increased contribution of more localised (fine-grained) patterns (Luppi et al., 2023b). This pattern was very comparable to classical psychedelics such as DMT, LSD, and psilocybin (Atasoy et al., 2017, 2018b; Vohryzek et al., 2024), even though they are believed to have different pharmacokinetics (Kadriu et al., 2021). In line with earlier theoretical predictions (Atasoy et al., 2018b), Luppi and colleagues (2023b) concluded that CHD can identify similar alterations in consciousness induced by different pharmacological interventions, both on the anaesthetic and psychedelic level.

Here, we examined which connectome harmonics – if any – exhibit heightened activity under the influence of ketamine-induced unresponsiveness as compared to wakefulness. To achieve this, we analysed frequency-specific changes in brain dynamics induced by ketamine by initially discretising the connectome-harmonic spectrum into 15 levels of wavenumbers k in the log-space. Then, we analysed the energy alterations within each bin of the harmonic spectrum for each of the conditions and for each subject individually. An overview of this procedure can be found in Figure 2, and a detailed account in the ‘Methods’ section.

**Figure 2.**
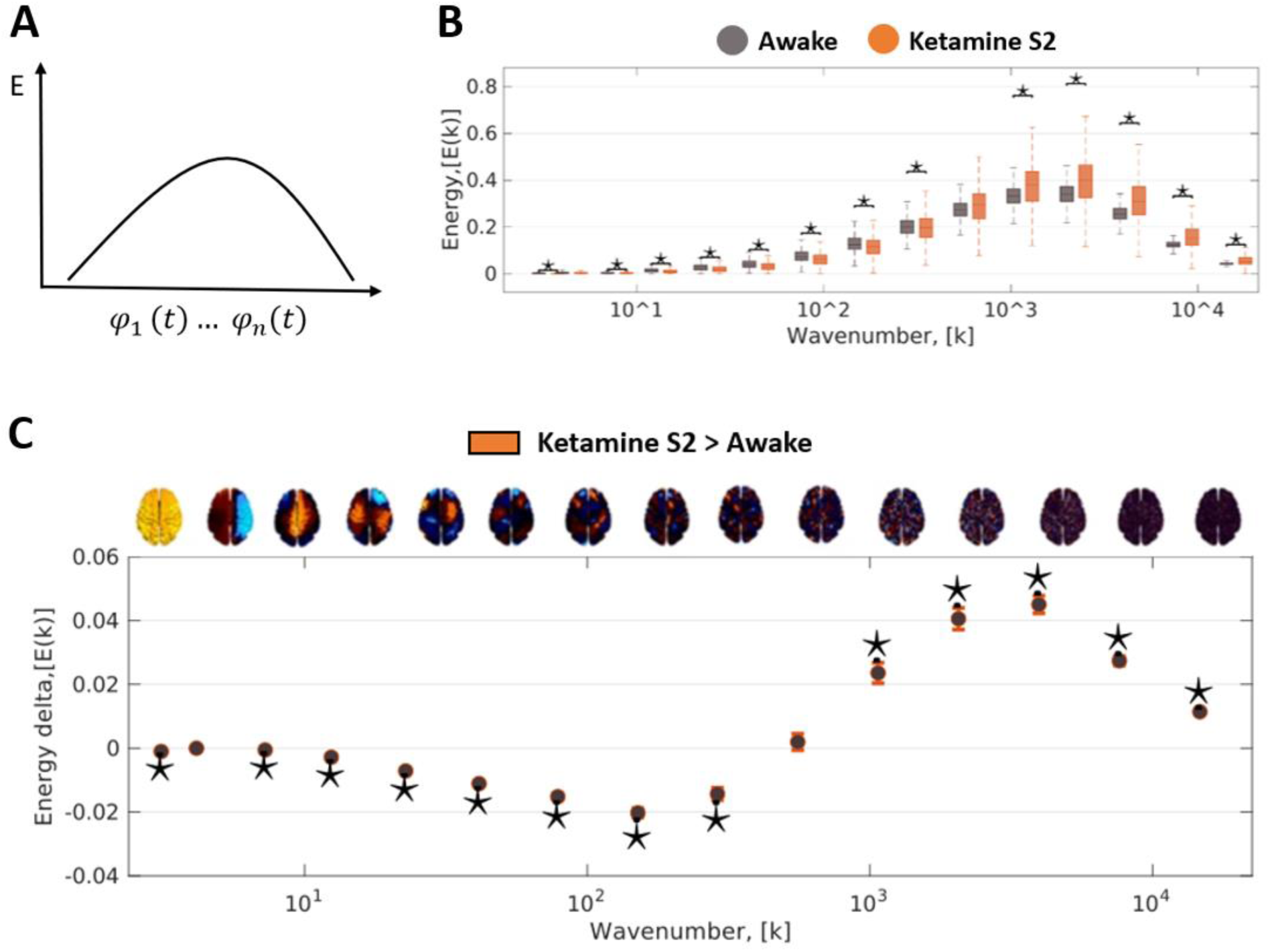
Connectome Harmonic Signature of Ketamine-Induced Unresponsiveness. **A** The connectome harmonic energy spectrum is determined by computing the square of the absolute contribution |*ω_k_*(*t*)|of individual harmonics to the fMRI data, which is then weighted by the square of the corresponding eigenvalue *λ*_k_ (representing intrinsic energy) for each timepoint *t ∈* {*1, …, T*}: *E*(*φ_k_, t*) = |*ω_k_*(*t*)|² *λ*_k_². **B** The comprehensive binned energy spectrum across subjects and timepoints is formed by discretising the energy of connectome harmonics into 15 logarithmically spaced frequency-specific bins. In this case, the target state is “Ketamine S2” (dark orange) and the reference state is “Awake” (dark grey). This approach has been demonstrated in previous studies to effectively emphasise the connectome harmonic signatures associated with altered states of consciousness (Atasoy et al., 2016, 2017, 2023; Luppi et al., 2023b). **C** A pair of conditions (Ketamine S2 vs Awake) was contrasted using linear mixed effects (LME) modelling, treating condition as a fixed effect and subjects as random effects. Additionally, timepoints were included as random effects nested within subjects. The plot displays the statistical estimates for the contrast, with error bars representing the 95% confidence intervals derived from the LME model. The specific comparison here: ketamine deep sedation (S2; *N* = 8 subjects with 300 timepoints each) > awake (*N* = 8 subjects with 300 timepoints each). A brain surface projection illustrating the connectome harmonic pattern associated with each frequency bin is displayed above the respective bin. These projections are averaged over the constituent spatial frequencies within each bin. *\*p* < 0.05, FDR-corrected across 15 frequency bins.

When comparing ketamine deep sedation (S2) to normal wakefulness using Linear Mixed Effect (LME) modelling, significant changes were observed in the energy levels of nearly all quantised wavelengths (*p* < 0.05; Figure 2B). An increase was found in the connectome harmonic energy spectrum across a broad range of high-frequency harmonics (wave numbers larger than 200 out of 18,715 connectome harmonics), while the energy spectrum of connectome harmonics decreased with wave numbers smaller than 200 (Figure 2C). This pattern was also found for lighter ketamine sedation (S1) compared to wakefulness, and even between the two different doses (S1 and S2; Figure S2). In addition, both doses of ketamine showed notably greater diversity in the range of connectome harmonics compared to normal wakefulness (Figure S3). Taken together, these results indicate that ketamine sedation is characterised by a more localised organisation of brain activity as compared to wakefulness. We can also conclude that the contribution of localised harmonics seems to increase with higher concentrations of ketamine, at least up to a certain point.

### Relating Ketamine’s Harmonic Signatures to the Signatures of Other Altered States of Consciousness

Previous research has shown that it is possible to generalise connectome harmonic patterns across datasets, thereby establishing the harmonic signature of both unconsciousness and the psychedelic state (Luppi et al., 2023b). To achieve this, Partial Least Squares Discriminant Analysis (PLS-DA) was used to comprehensively consider the entire spectrum of connectome harmonic changes simultaneously. This data-driven technique allowed the researchers to extract multivariate patterns of connectome harmonic energy, referred to as multivariate signatures (MVS), that effectively differentiate between pairs of conditions (Luppi et al., 2023b). The first principal component derived from PLS-DA represents the most discriminative pattern in the data, distinguishing subjects belonging to different states of consciousness. Luppi and colleagues (2023b) showed that this approach revealed two mirror-reversed multivariate patterns characterising the loss of consciousness (propofol anaesthesia and disorders of consciousness) and the psychedelic state (LSD and sub-anaesthetic ketamine). The same approach was then also used to show that the connectome harmonic signature of the serotonergic psychedelic DMT matches those of LSD and sub-anaesthetic ketamine (Vohryzek et al., 2024).

In the current study, the connectome harmonic signature observed for ketamine sedation was mapped onto signatures of other altered states of consciousness (propofol sedation, disorders of consciousness, ketamine sub-anaesthesia, psychedelic state induced by LSD), previously used by Luppi et al. (2023b), which allowed us to investigate the similarity with these states. To measure the alignment of the harmonic signatures of ketamine sedation and propofol sedation, we calculated the dot-product between the MVS that best discriminates between wakefulness and propofol moderate sedation (Luppi et al., 2023b), and the MVS that best discriminates between wakefulness and ketamine deep sedation (S2). The same procedure was applied to compare the CHD signature of ketamine-induced unresponsiveness (S2) to that of disorders of consciousness (DOC), obtained from comparing DOC patients who could or could not respond to tasks in the fMRI scanner. Finally, we also compared the signature of ketamine deep sedation (S2) to that of ketamine sub-anaesthesia, and to the CHD signature of the psychedelic state induced by LSD.

The dot-product is a mathematical operation that measures the extent to which two vectors point in the same direction, giving a positive value for vectors that point in roughly the same direction and a negative value for vectors that point in roughly the opposite direction. In the current study, a positive result indicates an alignment between two different MVSs associated with different states of consciousness, while a negative result indicates misalignment.

As can be seen in Figure 3, the results showed that the connectome harmonic signature of ketamine anaesthesia aligns with the signatures seen under the influence of LSD (Δ LSD-MVS projection = 0.0113, *p* < 0.001) and a sub-anaesthetic dose of ketamine (Δ ketamine-MVS projection = 0.0087, *p* < 0.001), as identified by Luppi and colleagues (2023b). On the other hand, the connectome harmonic signature of ketamine anaesthesia was inversely related to the signature seen under the influence of propofol anaesthesia (Δ propofol-MVS projection = −0.0213, *p* < 0.001) and the signature of patients with DOC (Δ DOC-MVS projection = 0.0206, *p* < 0.001). Similar results were found for a lighter sedative dose of ketamine (S1; Figure S4). This confirms earlier remarks that ketamine sedation, like atypical (ketamine sub-anaesthesia) and typical (LSD) psychedelics, gives rise to a more localised organisation of brain activity. This pattern is reversed in propofol sedation and in patients with DOC (Luppi et al., 2023b).

**Figure 3.**
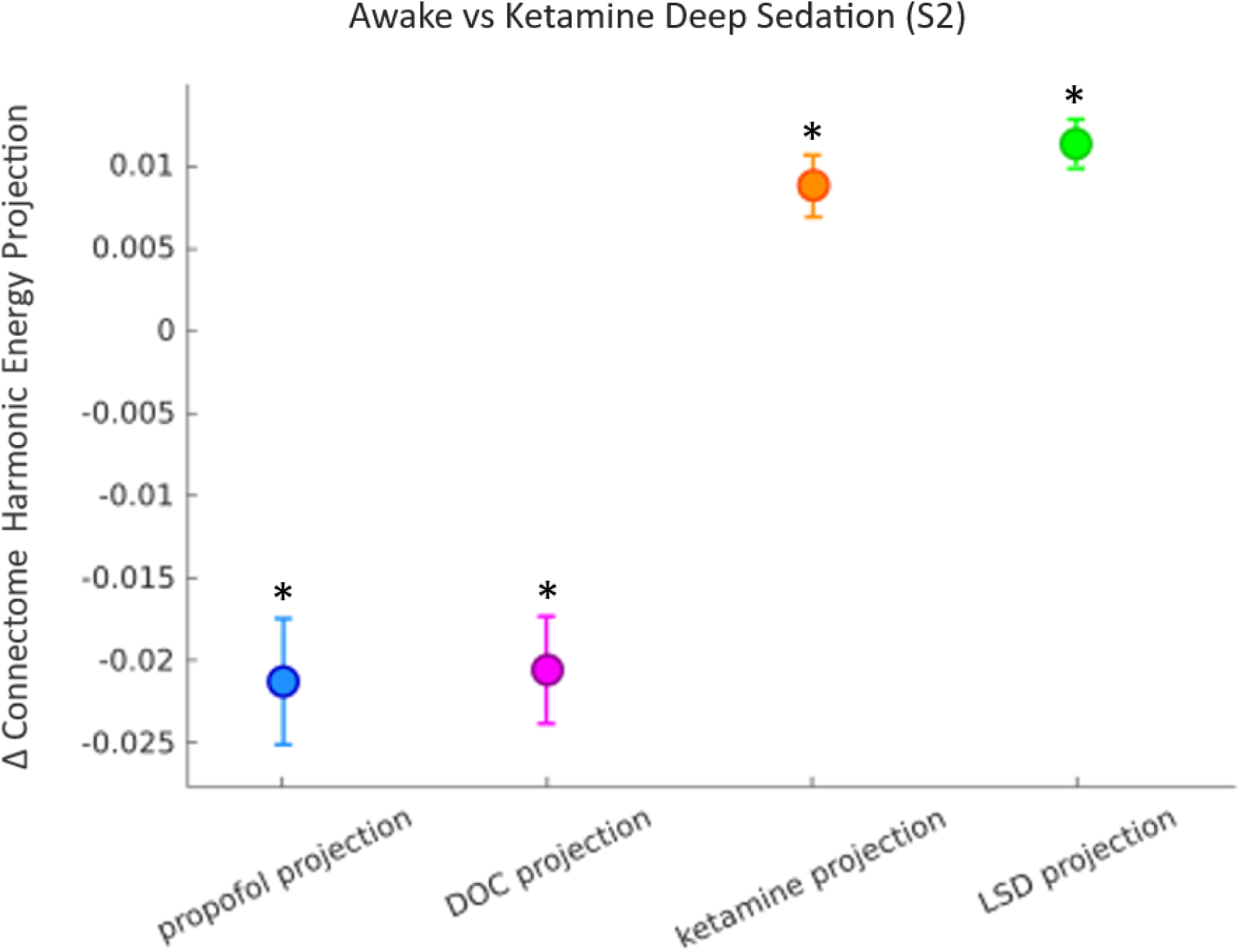
The Connectome Harmonic Signature of Ketamine-Induced Unresponsiveness Aligns More With the Signatures of Psychedelic States. The figure displays the drug-induced change in alignment (dot-product) between the spectrum of connectome harmonic energy of one state, and the multivariate energy signature (MVS) of another state. The dot-product between the MVS that best discriminates between “Awake” and “Propofol Moderate” (Moderate minus Awake; Luppi et al., 2023b), and the MVS that best discriminates between “Awake” and “Ketamine S2” is shown in blue. The dot-product between the MVS that best discriminates between DOC patients who could not respond to tasks in the fMRI scanner and DOC patients who could (fMRI-minus fMRI+), and the MVS that best discriminates between “Awake” and “Ketamine S2” is shown in violet. The dot-product between the MVS that best discriminates between “Ketamine Sub-anaesthesia” and “Placebo” (Ketamine minus Placebo), and the MVS that best discriminates between “Awake” and “Ketamine S2” is shown in orange. The dot-product between the MVS that best discriminates between “LSD” and “Placebo” (LSD minus Placebo), and the MVS that best discriminates between “Awake” and “Ketamine S2” is shown in green. The projections for propofol, DOC, ketamine, and LSD are obtained with permission of Luppi et al. (2023b). The points denote the means of the alignment (dot-product) of the MVS signatures across participants and the error bars indicate the standard errors of this alignment. *\*p* < 0.001

### Are the Results Replicable with a Different Human Structural Connectome?

Previous research has demonstrated the test-retest reliability of CHD using two scans of the same individuals during resting wakefulness. The evidence indicated that connectome harmonic signatures remain stable across scans of the same individuals, when they are in the same state of consciousness (Luppi et al., 2023b). This provides an important control for the results described previously.

Crucially, the results presented in this report are not explained by motion artifacts of the participants (Figure S7) and do not depend on the specific operationalisation of the human connectome. After employing an alternative method to reconstruct a high-resolution representation of the human connectome – combining a much larger sample of 985 Human Connectome Project (HCP) subjects, corresponding to a 100-fold increase in sample size – the results indicate that there is not much difference between the small and large HCP sample (Figure S5, S6), making connectome harmonics an especially suitable framework for mapping the landscape of consciousness across individuals and datasets.

## Discussion

The aim of this study was to investigate the organisation of brain activity under ketamine-induced unresponsiveness using the recently developed framework of connectome harmonic decomposition (CHD; Atasoy et al., 2016, 2017). Like the Fourier transform reflecting fundamental frequencies in temporal brain signals, the CHD method explicitly represents brain activity as fundamental harmonic modes by considering contributions from various spatial scales of the underlying structural network. Remember that the transition from low-to high-frequency connectome harmonics indicates a shift from global (coarse-grained) to more localised (fine-grained) activity patterns. Hence, CHD is in line with literature that considers brain-wide functional organisation, unlike isolated spatial regions in the brain, to be informative of consciousness (Adapa et al., 2013; Bonhomme et al., 2016; Boveroux et al., 2010; Craig et al., 2020; Demertzi et al., 2015, 2019; Di Perri et al., 2016, 2018; Guldenmund et al., 2016; Hannawi et al., 2015; Kafashan et al., 2016; Luppi et al., 2019; MacDonald et al., 2015; Palanca et al., 2015; Ranft et al., 2016; Spindler et al., 2021; Stamatakis et al., 2010; Threlkeld et al., 2018). In addition, CHD seems to reliably track alterations of consciousness across various datasets of healthy individuals and DOC patients, and across various pharmacological compounds (Atasoy et al., 2017, 2018b; Luppi et al., 2023b). Finally, the connectome harmonic signature obtained by CHD seems to be responsive to variations in dosage, as well as behaviourally indistinguishable subgroups of patients with DOC (Luppi et al., 2023b).

Recently, a study by Pang and colleagues (2023) observed that “structural eigenmodes (harmonic modes) derived solely from the brain’s geometry provide a more compact, accurate and parsimonious representation of its macroscale activity than alternative connectome-based models.” However, Faskowitz and colleagues (2023) in a commentary on this work argued that this claim contradicts prevailing theories regarding information flow in the brain, highlighting the significance of long-range axonal connections and bundled white matter in facilitating signal relay among cortical areas (Honey et al., 2009; Deco et al., 2011; Seguin et al., 2023). Despite these controversies, a recent study was able to demonstrate that, connected through the Exponential Distance Rule (EDR), cortical short-range connectivity and geometry are two sides of the same coin: being spatially embedded, structural connections implicitly encode geometry, with an additional important role of long-range connectivity exceptions to the EDR. Thus, according to Vohryzek, Kringelbach and Deco (2024), the best results of explaining functional activity patterns in terms of underlying eigenmodes seem to be obtained by incorporating both short-range connectivity via the EDR and long-range connectivity embodied by the brain’s white-matter tracts, adding merit to the framework used in the current study.

Our results show that ketamine-induced unresponsiveness increases the contribution of high-frequency (fine-grained) connectome harmonics, and decreases the contribution of low-frequency (coarse-grained) connectome harmonics. As described previously, others have interpreted increased prevalence of low-frequency harmonics as reflecting activity that is more structurally-coupled, and high-frequency harmonics as representing structurally decoupled activity (Preti & Van de Ville, 2019). In this context, our findings would correspond to structure-function decoupling of brain activity in ketamine-induced unresponsiveness, in accordance with previous observations with classic psychedelics.

A substantial body of evidence has shown that this shift from low-to high-frequency harmonics is reliably associated with an altered state of consciousness seen in people under the influence of serotonergic psychedelic compounds (psilocybin, LSD, and DMT; Atasoy et al., 2017, 2018b; Vohryzek et al., 2024), sub-anaesthetic doses of ketamine (Luppi et al., 2023b), and in meditation (Atasoy et al., 2023). Although this seems contradictory to the fact that the volunteers in this study were behaviourally unresponsive, the results could potentially be explained by their anecdotal reports of “psychedelic-like” dreams during ketamine sedation (Bonhomme et al., 2016), indicating that some form of consciousness might have been preserved. These dreams encompassed sensations of well-being, joy, and peace, along with perceptions of an otherworldly environment, flying, encountering a bright light, experiencing daydream-like states, feeling dissolved into the surroundings, body distortions, sensations of falling, feelings of confinement, and the perception of impending death (Bonhomme et al., 2016). So, although volunteers are usually completely unresponsive when given high doses of ketamine, some sort of conscious experience seems to be preserved, as they are able to report afterwards. This is in line with previous studies investigating the occurrence of dreams and sensations in ketamine deep sedation (Hejja & Galloon, 1975; Jansen, 2001; Maya et al., 2018; Sarasso et al., 2015).

Furthermore, our findings seem to map well onto the current neuroscientific literature investigating ketamine sedation. For example, electroencephalogram (EEG) studies have shown that ketamine-induced unresponsiveness maintains spatiotemporal complexity, as measured by the Perturbational Complexity Index (PCI; Sarasso et al., 2015), as compared to propofol, midazolam and xenon anaesthesia, which all seem to show decreased PCI (Casali et al., 2013). More detailed analyses of the dose-dependent effects of ketamine on EEG signals have shown a more nuanced story, indicating that the spatiotemporal complexity linked to ketamine-induced state changes exhibits characteristics resembling those of general anaesthesia (low complexity), altered states of consciousness (high complexity), and normal wakefulness (Farnes et al., 2020; Li & Mashour, 2019). Another study confirmed these results, showing that ketamine deep sedation was found to show dynamic shifts in the organisation of brain states, transitioning from a diminished repertoire richness to a level comparable to that observed during normal wakefulness right before recovery (Li et al., 2022). This is consistent with a disruption of connected consciousness during ketamine deep sedation, marked by vivid experiences that are disconnected from the environment (e.g., dreams, hallucinations), alongside a complete unawareness of environmental events (Grace, 2003).

In the current study, we also showed that the connectome harmonic signatures seen in ketamine-induced unresponsiveness are more aligned with those seen in sub-anaesthetic ketamine and classical serotonergic psychedelics than those seen in propofol sedation and DOC. The overlap between ketamine and classical serotonergic psychedelics (e.g., LSD, psilocybin) has been identified by previous research. Although operating on distinct pharmacological mechanisms (Kadriu et al., 2021), both ketamine and classical psychedelics promote alterations in neuroplasticity, including the growth of neurites, the formation of synapses, and the reinforcement of synaptic connections (Lima da Cruz et al., 2018; Ly et al., 2018; Moda-Sava et al., 2019; Phoumthipphavong et al., 2016; Treccani et al., 2019). In addition, ketamine and serotonergic psychedelics both induce similar phenomenology (Studerus, Gamma & Vollenweider, 2010), have antidepressant effects (Berman et al., 2010; Galvão-Coelho et al., 2021; Husain et al., 2022; Xu et al., 2022), increase neurophysiologic complexity (Farnes et al., 2020; Li & Mashour, 2019; Schartner et al., 2017; Dai et al., 2023), and expand the repertoire of functional brain connectivity modes (Li et al., 2022; Luppi et al., 2023b). A recent study by Luppi and colleagues (2023a) used in-vivo maps of neurotransmitter distribution in the human brain obtained from Positron Emission Tomography to examine the macroscale neurotransmitter signatures of a wide range of pharmacological agents. They found that ketamine-induced unresponsiveness showed substantial similarities with serotonergic psychedelics such as LSD, as well as sub-anaesthetic ketamine. Together with the current investigation, we conclude that the neurotransmitter profiles and connectome harmonic signatures corresponding to ketamine deep sedation align with molecular and subjective effects, rather than with behavioural effects.

Our findings also confirm the crucial role played by fine-grained, high-frequency harmonics in distinguishing between different states of consciousness (Atasoy et al., 2017; Luppi et al., 2023b). This valuable insight was made possible by leveraging high-quality, high-resolution diffusion data from the Human Connectome Project (HCP), allowing to achieve resolutions up to three orders of magnitude finer than other methods of harmonic mode decomposition relying on parcellated data (Abdelnour et al., 2014; Glomb et al., 2020; Medaglia et al., 2018; Preti & Van De Ville, 2019; Wang et al., 2017; Xie et al., 2021). In addition, the results were successfully replicated using a high-resolution connectome constructed from data collected from 985 HCP subjects, representing one of the most comprehensive characterisations of the structural connectivity of the human brain to date (Luppi et al., 2023b), and demonstrating that our results are not just due to the specific implementation of the human structural connectome used.

However, it is important to acknowledge that obtaining high-quality connectome reconstructions may present challenges (Sarwar, Ramamohanarao & Zalesky, 2021). First, variability in data quality, arising from factors such as imaging artifacts, motion during scanning, and signal-to-noise ratio, can introduce inaccuracies in the reconstructed connectome (Setsompop et al., 2012; Toga & Thompson, 2003). Second, the human brain exhibits significant inter-subject variability in terms of anatomy, connectivity patterns, and functional organisation. Accounting for this variability and establishing reliable norms for connectome reconstruction across individuals is challenging (Van Essen & Ugurbil, 2012). Third, combining data from multiple imaging modalities, such as structural MRI, diffusion MRI, and functional MRI, poses challenges in terms of data fusion and integration. Aligning data from different modalities to construct a comprehensive connectome requires sophisticated processing and validation methods (Jeurissen et al., 2019). Fourth, diffusion imaging and tractography are unable to accurately infer fibre directionality, which is a crucial aspect of brain wiring (Schilling et al., 2019). Taking into account these limitations, the connectome used in this study stands out as one of the most thorough descriptions of the structure within the human brain to date, adding credibility to our results.

Another limitation of this study is the rather low number of participants (*N* = 8). With a smaller sample size, the statistical power of the study decreases, making it more difficult to detect true effects or relationships between variables, leading to an increased likelihood of false-negative results (Button et al., 2013). Furthermore, findings with a small number of participants may not generalise well to other individuals or groups (Szucs & Ioannidis, 2017). Nevertheless, the fact that the obtained results in this study prove robust (e.g., *p*-values well below the significance threshold), were generalised across different connectomes, and align with previous investigations, may mitigate some of these shortcomings.

Given that our investigation confirms the previous conclusion that CHD seems to reliably track consciousness in the absence of behavioural responsiveness, it could serve as a valuable tool in clinical settings to identify patients with DOC who warrant further assessment. This screening tool could help reduce the rate of misdiagnosis observed in these DOC patients when relying solely on behavioural criteria (Luppi et al., 2021a; Naci et al., 2017). Replicating current findings in diverse and larger samples of both DOC patients and individuals undergoing pharmacological perturbations of consciousness is essential to validate their reliability and robustness. Also, to make our findings clinically relevant, a more individualised approach will need to be taken (Laforge et al., 2020).

In conclusion, the robust analytical approach of CHD was used to investigate the brain dynamics of ketamine-induced unresponsiveness, unveiling fresh insights into both the neural underpinnings of this unique state of anaesthesia and its relationship to human consciousness. The findings underscore the importance of considering global brain function and associated subjective experiences in terms of the dynamic activation of harmonic brain states (connectome harmonics). Notably, this perspective reveals the dynamic spectrum of brain activity and suggests a shift from a global organisation of brain activity during wakefulness to more localised activity patterns during ketamine-induced unresponsiveness. This particular signature maps well unto the signatures seen in classical psychedelics (DMT, LSD, and psilocybin) and subanaesthetic ketamine, which can be related to the “psychedelic-like” dream reports of the volunteers. Overall, the results align with the previous hypothesis that the connectome harmonic decomposition (CHD) analysis could potentially be used to track consciousness in participants who, from the outside, are completely unconscious when assessed based on behavioural responsiveness.

## Data Availability

The raw ketamine data are available upon request from author VB.

## Acknowledgements

Canadian Institute for Advanced Research (CIFAR; grant RCZB/072 RG93193) [to EAS];

Stephen Erskine Fellowship at Queens’ College, Cambridge [to EAS];

Belgian National Funds for Scientific Research (F.R.S-FNRS) [to PC and NLNA];

GIGA-Doctoral School for Health Sciences (University of Liège) [to PC];

Human Brain Project [to NLNA];

EU H2020 Future and Emerging Technologies (FET) Proactive project Neurotwin grant agreement no. 101017716 [to JV]

European Research Council Consolidator Grant: CAREGIVING (615539), Pettit Foundation, Carlsberg Foundation and Center for Music in the Brain, funded by the Danish National Research Foundation (DNRF117) [to MLK].

Engineering and Physical Sciences Research Council (capital grant EP/T022159/1),

DiRAC funding from the Science and Technology Facilities Council;

MRC research infrastructure award (MR/M009041/1).

