## Supplementary Figures for "Ketamine-Induced Unresponsiveness Shows a Harmonic Shift from Global to Localised Functional Organisation"

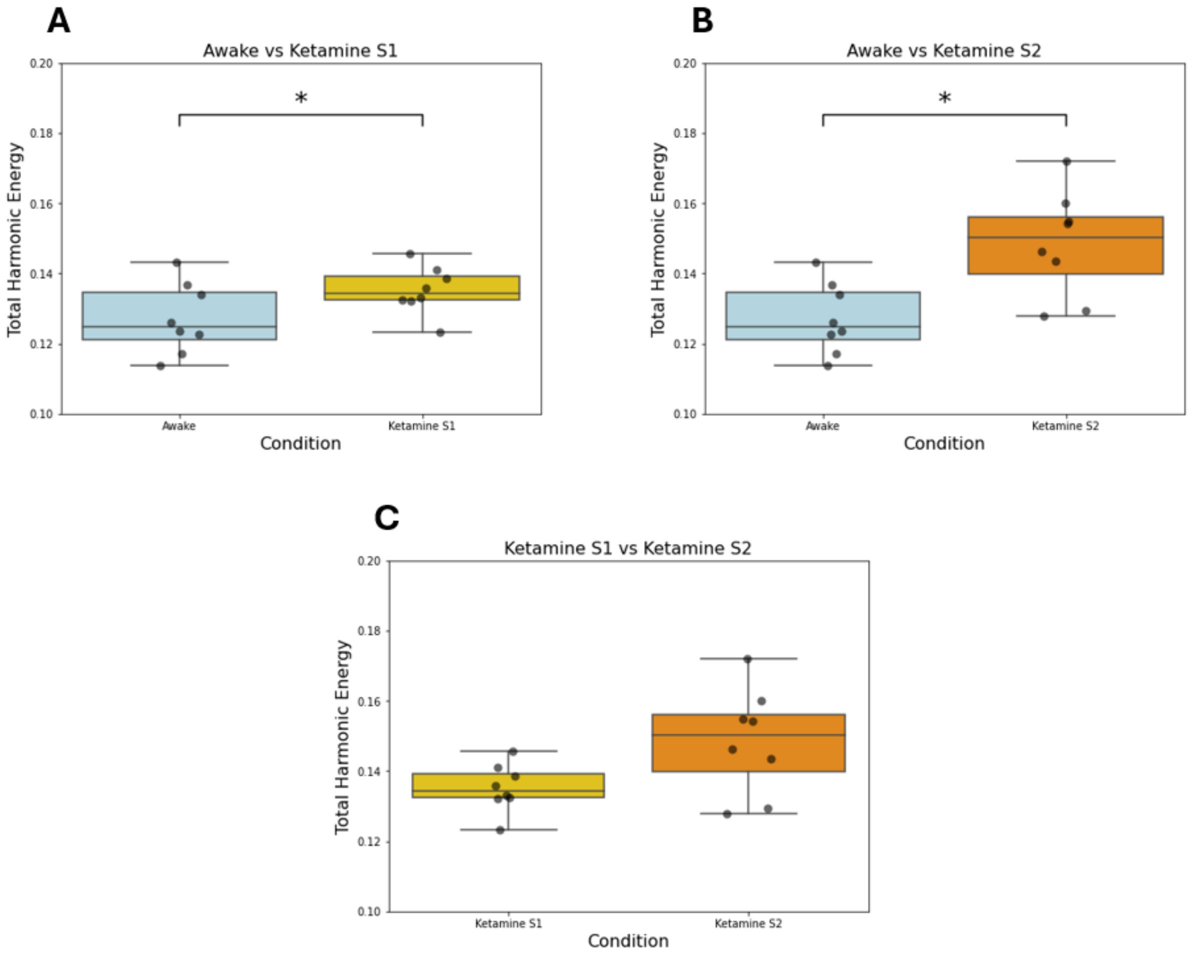


**Figure S1. Changes in total energy of brain states under the influence of ketamine sedation.**

**A** Total energy of all connectome harmonics in a state of wakefulness (Awake) and under the influence of ketamine sedation (Ketamine S1). The observed difference between the two conditions was significant ($p = 0.0257$, repeated-measures t-test). **B** Total energy of all connectome harmonics in a state of wakefulness (Awake) and under the influence of ketamine deep sedation (Ketamine S2). The observed difference between the two conditions was significant ($p = 0.0146$, repeated-measures t-test). **C** Total energy of all connectome harmonics under two different doses of ketamine sedation. The observed difference between the conditions was not significant ($p = 0.0788$, repeated-measures t-test).

* $p < 0.05$

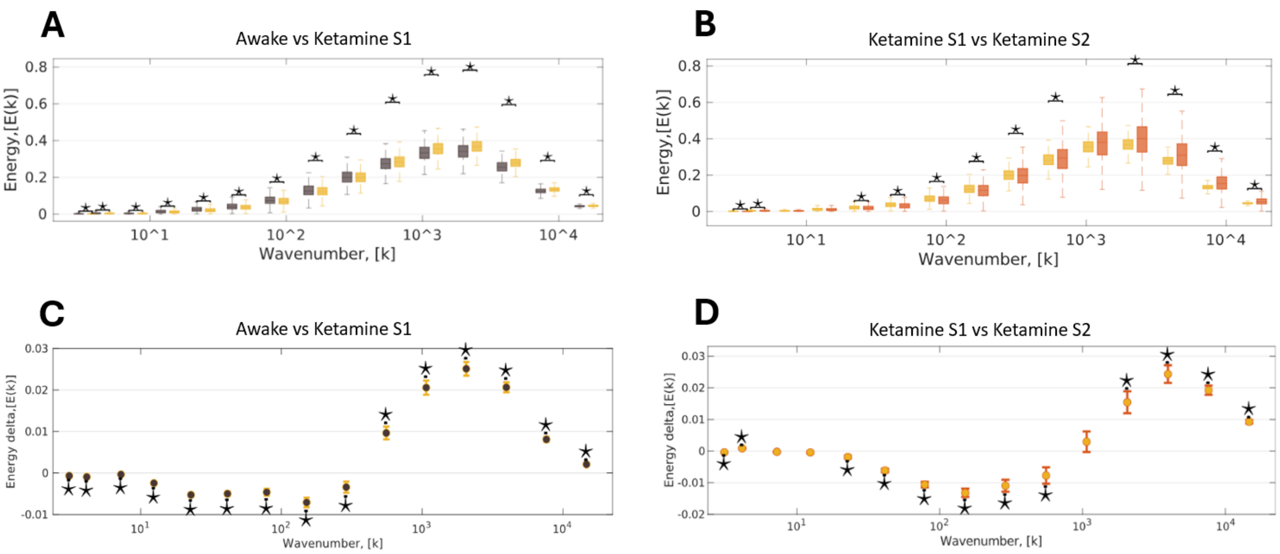


**Figure S2. Connectome harmonic energy signature of ketamine anaesthesia.**

**A** The binned energy spectrum across subjects and timepoints ($N = 8$, 300 timepoints each), with ketamine sedation (Ketamine S1) as target state and wakefulness (Awake) as reference state. **B** Statistical estimates from LME modelling between wakefulness (Awake) and ketamine sedation (Ketamine S1), treating condition as a fixed effect and subjects as random effects. Timepoints were also incorporated as random effects, nested within subjects. **C** The binned energy spectrum across subjects and timepoints ($N = 8$, 300 timepoints each), with “Ketamine S2” as target state and “Ketamine S1” as reference state. **D** Statistical estimates from LME modelling between “Ketamine S1” and “Ketamine S2”, treating condition as a fixed effect and subjects as random effects. Timepoints were also incorporated as random effects, nested within subjects.

Characteristics of the boxplots are as follows: central line indicates the median, box edges represent 25$^{th}$ and 75$^{th}$ percentiles, whiskers resemble 1.5x inter-quartile interval. This is consistent with the data presented in Luppi et al. (2023). *$p < 0.05$, FDR-corrected across 15 frequency bins.

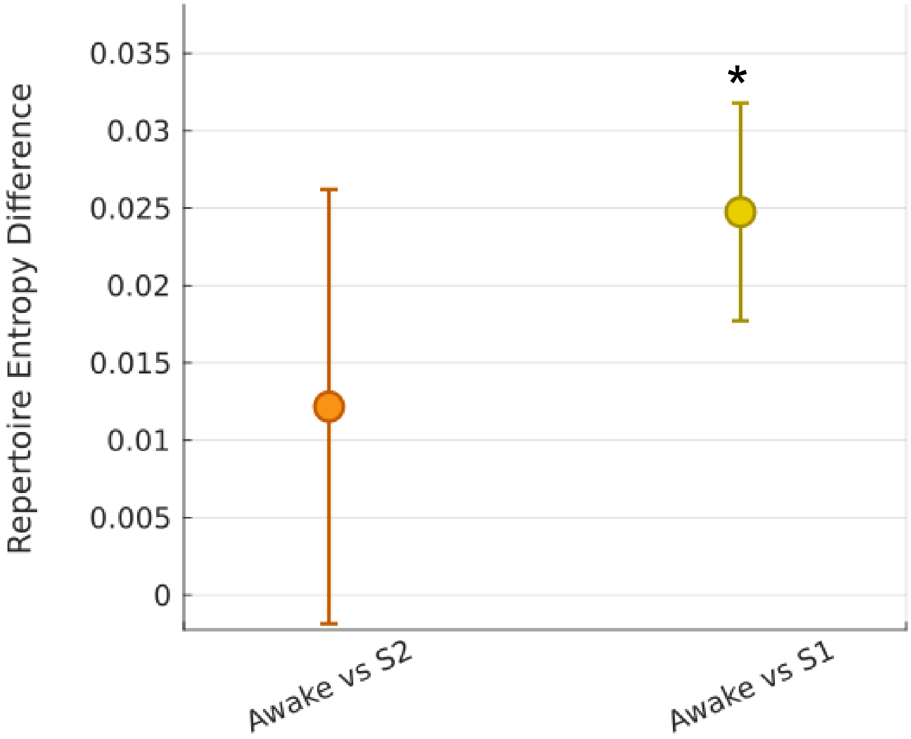


**Figure S3. Diversity of connectome harmonic repertoire of ketamine anaesthesia.**

Pairs of conditions (Awake vs Ketamine S2, Awake vs Ketamine S1) were compared using linear mixed effects modelling, treating condition as a fixed effect and subjects as random effects. Timepoints were also included as random effects nested within subjects. The plot displays the statistical estimates for each contrast, with error bars representing the 95% confidence intervals from the LME model. At each timepoint, the contribution of each connectome harmonic to the overall brain activity pattern is measured by its power $P(\varphi_k, t) = |\omega_k(t)|$. The diversity of the harmonic power repertoire is quantified by the entropy of the power distribution, with a higher repertoire representing a broader range of connectome harmonics involved in cortical activity composition. The difference in repertoire entropy between "Awake" and "Ketamine S2" was not significant ($p = 0.0889$, repeated-measures t-test), while the difference between "Awake" and "Ketamine S1" was significant ($p = 6.0484e^{-12}$, repeated-measures t-test).

* $p < 0.001$

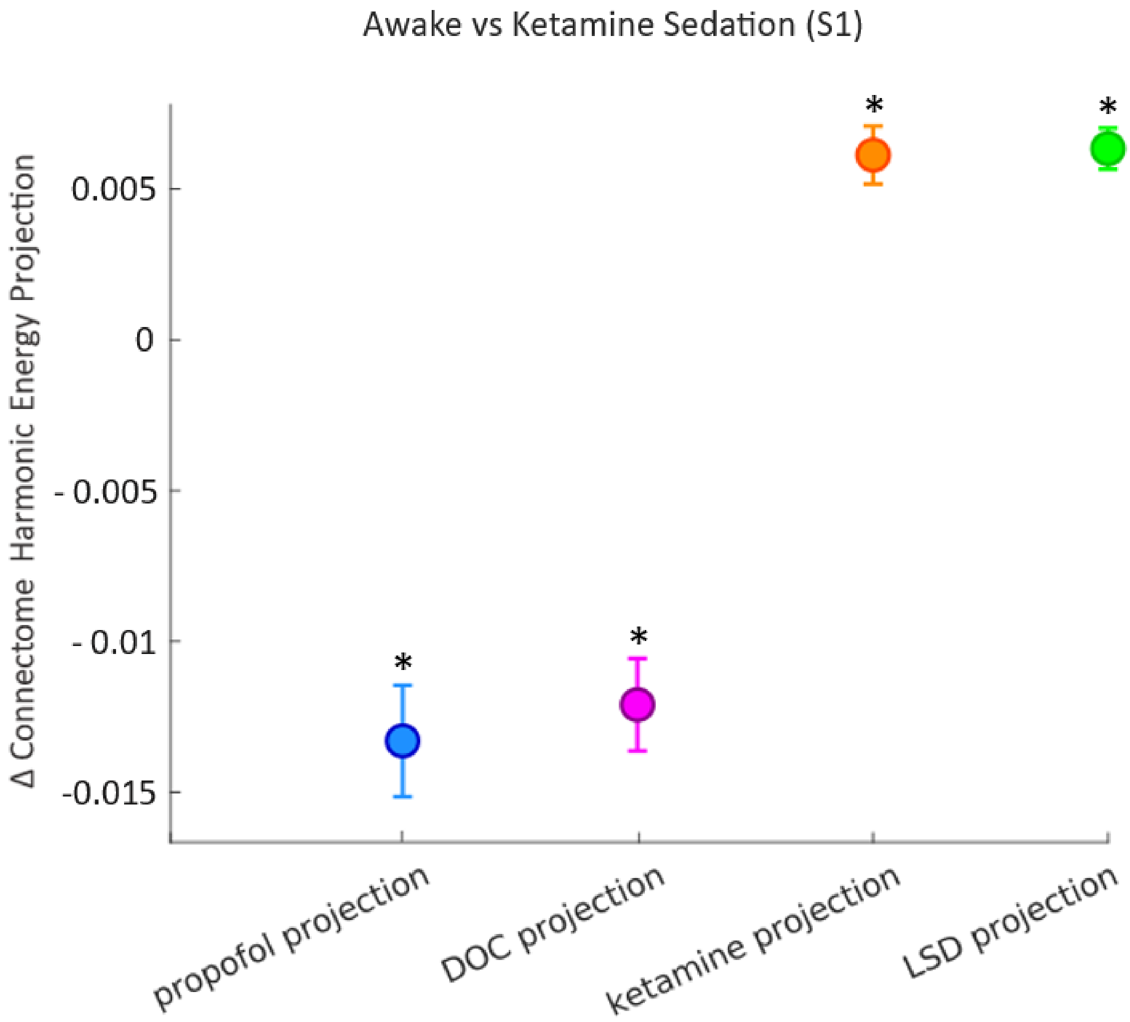


**Figure S4. Alignment of PLS-DA First component loadings for ketamine S1.**

The figure displays the drug-induced change in alignment (dot-product) between the spectrum of connectome harmonic energy of one state, and the multivariate energy signature (MVS) of another state. The dot-product between the MVS that best discriminates between “Awake” and “Propofol Moderate” (Moderate minus Awake; Luppi et al., 2023b), and the MVS that best discriminates between “Awake” and “Ketamine S1” is shown in blue. The dot-product between the MVS that best discriminates between “fMRI-” and “fMRI+” (fMRI- minus fMRI+), and the MVS that best discriminates between “Awake” and “Ketamine S1” is shown in violet. The dot-product between the MVS that best discriminates between “Ketamine Sub-anaesthesia” and “Placebo” (Ketamine minus Placebo), and the MVS that best discriminates between “Awake” and “Ketamine S1” is shown in orange. The dot-product between the MVS that best discriminates between “LSD” and “Placebo” (LSD minus Placebo), and the MVS that best discriminates between “Awake” and “Ketamine S1” is shown in green. The projections for propofol, DOC, ketamine, and LSD are obtained with permission of Luppi et al. (2023b).

The points denote the means of the alignment (dot-product) of the MVS signatures across participants and the error bars indicate the standard errors of this alignment.

*$p < 0.001$

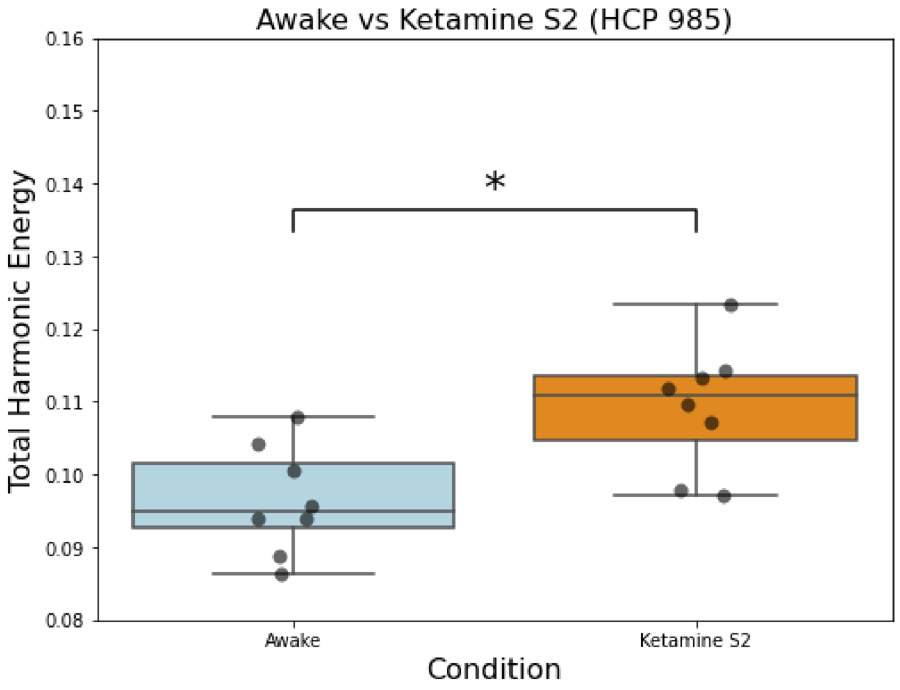


**Figure S5. Changes in total energy of brain states under the influence of an anaesthetic dose of ketamine, for the HCP-985 connectome.**

Total energy of all connectome harmonics in a state of wakefulness (Awake) and under the influence of ketamine deep sedation (Ketamine S2). The observed difference between the two conditions was significant ($p$ = 0.0306, repeated-measures t-test).

* $p < 0.05$

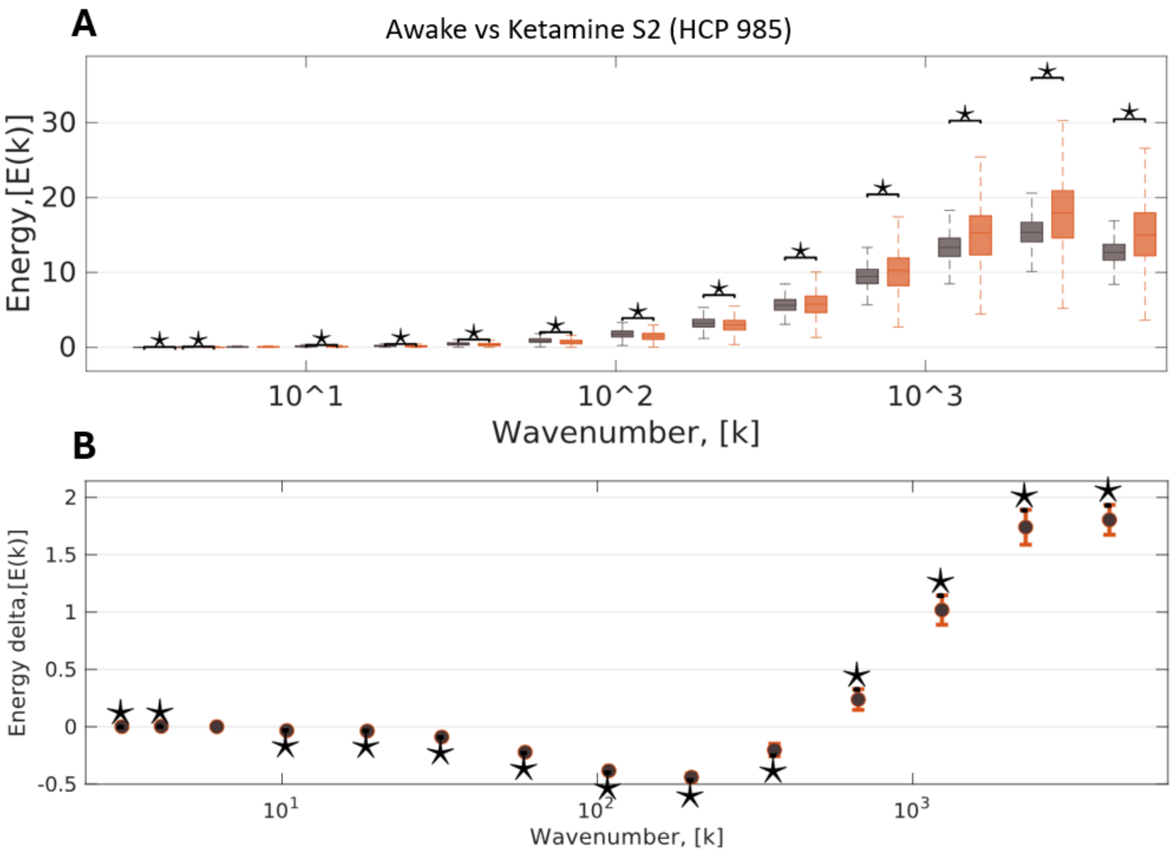


**Figure S6. Connectome harmonic energy signature of ketamine-induced unresponsiveness (S2), for the HCP-985 connectome.**

**A** The binned energy spectrum across subjects and timepoints ($N$ = 8, 300 timepoints each), with ketamine deep sedation (Ketamine S2) as target state and wakefulness (Awake) as reference state. **B** Statistical estimates from LME modelling between wakefulness (Awake) and ketamine deep sedation (Ketamine S2), treating condition as a fixed effect and subjects as random effects. Timepoints were also incorporated as random effects, nested within subjects.

Characteristics of the boxplots are as follows: central line indicates the median, box edges represent 25th and 75th percentiles, whiskers resemble 1.5x inter-quartile interval. This is consistent with the data presented in Luppi et al. (2023). *$p < 0.05$, FDR-corrected across 15 frequency bins.

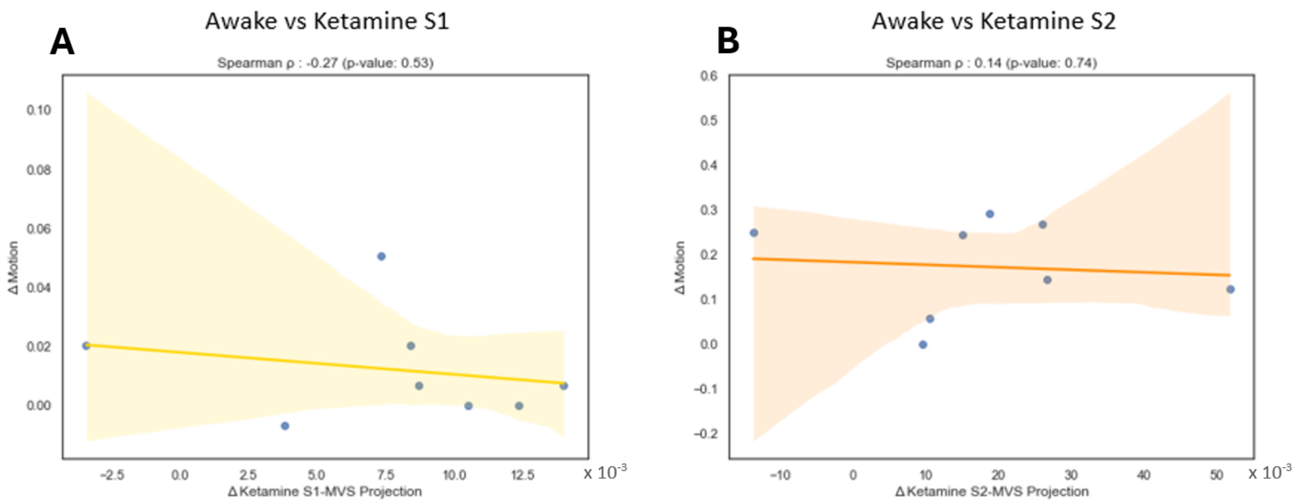


**Figure S7. Projection of individual connectome harmonic signatures onto multivariate energy signature (MVS) does not significantly correlate with differences in subject motion in the scanner.**

**A** Delta in ketamine (S1) connectome harmonic energy projection onto the MVS discriminating between “Awake” and “Ketamine S1”, versus the delta in head motion (Ketamine S1 minus Awake; $N = 8$, $\rho = -0.27$, $p = 0.53$). **B** Delta in ketamine (S2) connectome harmonic energy projection onto the MVS discriminating between “Awake” and “Ketamine S2”, versus the delta in head motion (Ketamine S2 minus Awake; $N = 8$, $\rho = 0.14$, $p = 0.74$).
